# scProfiterole: Clustering of Single-Cell Proteomic Data Using Graph Contrastive Learning via Spectral Filters

**DOI:** 10.64898/2026.02.26.708196

**Authors:** Mustafa Coşkun, Filipa Blasco Lopes, Pınar Kubilay Tolunay, Mark R. Chance, Mehmet Koyutürk

## Abstract

Novel technologies for the acquisition of protein expression data at the single cell level are emerging rapidly. Although there exists a substantial body of computational algorithms and tools for the analysis of single cell gene expression (scRNAseq) data, tools for even basic tasks such as clustering or cell type identification for single cell proteomic (scProteomics) data are relatively scarce. Adoption of algorithms that have been developed for scRNAseq into scProteomics is challenged by the larger number of drop-outs, missing data, and noise in single cell proteomic data. Graph contrastive learning (GCL) on cell-to-cell similarity graphs derived from single cell protein expression profiles show promise in cell type identification. However, missing edges and noise in the cell-to-cell similarity graph requires careful design of convolution matrices to overcome the imperfections in these graphs. Here, we introduce scProfiterole (Single Cell Proteomics Clustering via Spectral Filters), a computational framework to facilitate effective use of spectral graph filters in GCL-based clustering of single cell proteomic data. Since clustering assumes a homophilic network topology, we consider three types of homophilic filters: (i) random walks, (ii) heat kernels, (iii) beta kernels. Direct implementation of these filters is computationally prohibitive, thus the filters are either truncated or approximated in practice. To overcome this limitation, scProfiterole uses Arnoldi orthonormalization to implement polynomial interpolations of any given spectral graph filter. Our results on comprehensive single cell proteomic data show that (i) graph contrastive learning with learnable polynomial coefficients that are carefully initialized improves the effectiveness and robustness of cell type identification, (ii) heat kernels and beta kernels improve clustering performance over adjacency matrices or random walks, and (iii) polynomial interpolation of spectral filters outperforms approximation or truncation. The source code for scProfiterole and Supplementary Materials are available at https://github.com/mustafaCoskunAgu/scProfiterole.

## 1 Introduction

Recent technological advances in single-cell RNA sequencing (scRNA-seq) have enabled monitoring of transcriptomic states at the resolution of individual cells [34]. Transcriptomic states provide useful information to characterize cellular heterogeneity [25], cell-to-cell interactions [24,38], and spatio-temporal dynamics of cell states [18]. However, mRNA expression exhibits limited correlation with protein abundance [28,29], thus provides limited information on the post-translational states of the cells [2]. Proteins serve as the primary actors of cellular function and signaling, acting through enzymatic activity, complex formation, and post-translational modifications. Due to the difficulties in the identification and quantification of peptide sequences and the transient nature of post-translational dynamics, single-cell proteomics has long been limited to monitoring the expression of a small set of selected proteins [1]. The emergence of recent technologies for proteome-wide single-cell proteomics (scProteomics) shed novel insights into cell states by directly measuring functional cellular processes [14].

As single cell proteomic data emerges, the analysis of this novel type of omic data presents distinct computational challenges. Technical limitations in sample preparation, isotopic labeling, and mass spectrometry (MS) acquisition often introduce substantial peptide quantification uncertainty, missing data, and batch effects [32]. These issues are accentuated in single-cell proteomics [13], rendering such basic tasks as clustering and cell type annotation highly challenging [23]. While it may be possible to improve cell type identification by improving the construction of cell-to-cell similarity matrices, sparsity-noise trade-off and the imbalance of available data between different proteins pose important limitations. Motivated by these considerations, we here focus on the task of clustering cell-to-cell similarity graphs derived from scProteomics data and develop algorithms to overcome the limited reliability of these graphs.

Recent studies demonstrate that graph-based deep learning, particularly graph contrastive learning (GCL), can denoise proteomic data and model uncertainty to a certain extent [23]. Graph neural networks have also demonstrated success in analyzing scRNAseq data [4,37]. However, the application of graph representation learning to scProteomics can be hampered by the *over-smoothing* of signals with increasing number of layers in the neural network [5,22]. Fig. 1 demonstrates the detrimental effect of oversmoothing on the clustering of scProteomics data via scPROTEIN [23], a GCL algorithm that uses Graph Convolutional Network (GCN) based encoders. This limitation restricts the ability to aggregate information from distant yet functionally related nodes, an essential aspect of modeling biological systems that often display long-range homophily [8]. Indeed, scPROTEIN uses a two-layer neural network, limiting the view of the GCN to shallow neighborhood aggregation.

**Fig. 1:**
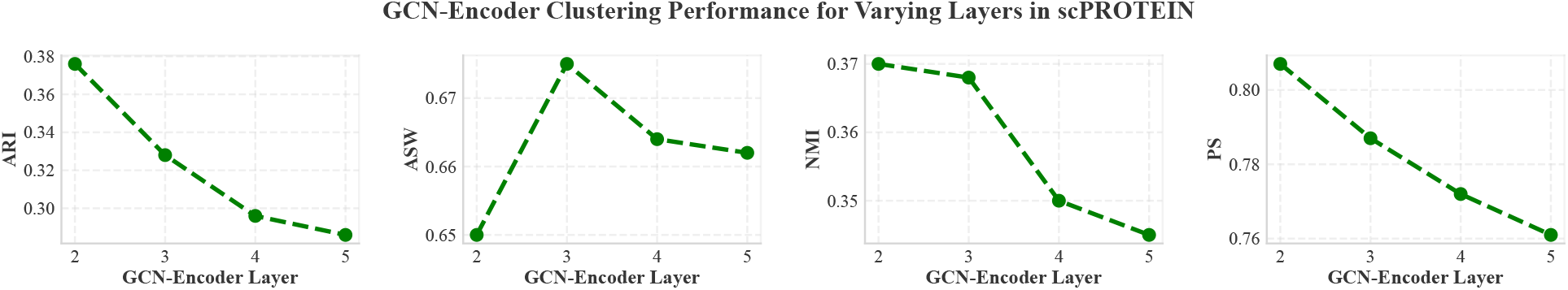
Performance of GCN encoders in clustering scProteomics data as a function of layer depth. The performance of encoders using the adjacency matrix of the cell-to-cell similarity matrix for convolution on the `Scope2_Specht` dataset is shown. Over-smoothing leads to performance degradation with deeper architectures. ARI: Adjusted Rand Index, ASW: Average Silhouette Width, NMI: Normalized Mutual Information, PS: Purity Score.

We argue that spectral graph filters can overcome the limitations posed by the sparse and noisy nature of cell-to-cell similarity graphs in scProteomics without requiring a deeper neural network, by providing a broader perspective of graph topology. It has been repeatedly shown in the graph machine learning literature [20], as well as in other applications of graph machine learning in systems biology [19], that spectral graph filters are highly effective in overcoming imperfections in graph-structured data. Spectral filters transform graph topology by effectively operating in the eigenspace of the graph adjacency matrix or the graph Laplacian, where the filter specifies the part of the spectra to be emphasized. In the eigen-decomposition of the (normalized) adjacency or Laplacian matrix of a graph, each eigenvalue-eigenvector pair represents a latent pattern in graph topology. Application of a spectral filter therefore corresponds to reweighing these latent patterns to compute a new matrix to be used in graph convolution, in which certain aspects of graph topology are amplified.

Since computing the eigen-decomposition of the filter functions is cost-prohibitive, these filters are usually implemented using polynomial representations. Importantly, implementation of the filter as a polynomial of the normalized adjacency matrix significantly enhances scalability. This is because, although the convolution matrix that results from spectral filtering can be dense, it never needs to be explicitly computed. Using the polynomial representation of the spectral filter, graph convolution can be implemented as a series of sparse matrix-vector multiplications, avoiding computation of eigen-decompositions or matrix-matrix multiplications. Furthermore, by designating the coefficients of the polynomial as learnable parameters, the filter can be adaptively learned during training. However, the learned coefficients and the accuracy of the model depend heavily on initialization [7], thus an important design choice is the selection of the spectral filter that is used to initialize these coefficients. Since cell-to-cell similarity graphs are constructed to be homophilic by design, low-pass filters stand as the natural choice for clustering these graphs.

Here, we present scProfiterole (Single Cell Proteomics Clustering via Spectral Filters), a framework that combines the interpretability of spectral graph theory with the power of graph contrastive learning. To provide a broad perspective on the use of low-pass filters in clustering single-cell proteomic data, scProfiterole implements three families of spectral filters: 1) Random-walk with restarts (RWR), 2) Heat kernels (HK), 3) Beta kernels (BK). These filters are visualized in Fig. 2. In the figure, the range of the eigenvalues of the normalized adjacency matrix of any undirected graph is shown on the x-axis. The y-axis shows the function that is applied to the eigenvalues of the graph, i.e, *g* (*ω* ) denotes the weight assigned to the latent pattern in graph topology that corresponds to eigenvalue *ω*. While RWRs are well-established and relatively straightforward to implement, the filters that correspond to RWRs focus on a narrow band of the eigenspectra and the damping factor (*α*) has limited effect on the filter (Fig. 2, left). Heat kernels, also called continuous-time random walks [21], offer a more flexible and broader family of filters (Fig. 2, right), but they are more difficult to implement. Taylor approximation of Heat kernels is used for efficient implementation [21]. Finally, Beta kernels (Fig. 2, center) offer a family of filters that are polynomial by design, thus they can be implemented directly.

**Fig. 2:**
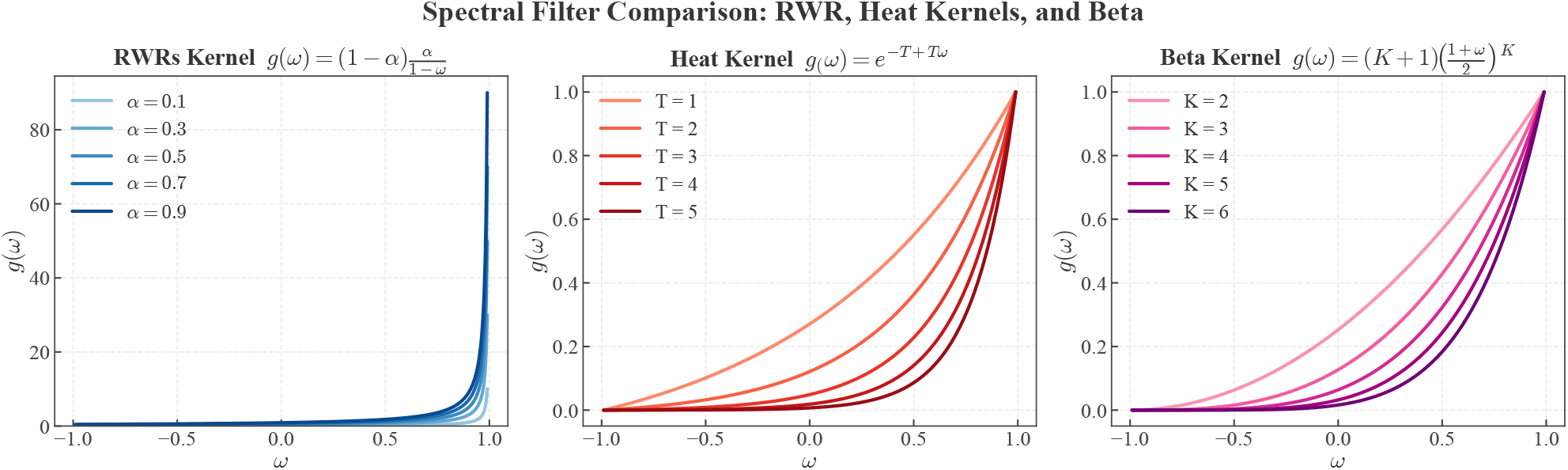
The landscape of spectral filters represented by the low-pass filter families implemented in scProfiterole. The x-axis shows the range of the eigenvalues of the normalized adjacency matrix of a graph, the plots show the function represented by the specific type of filter with the specific value of the filter’s parameter. The filters appear reversed (i.e, they are monotonically increasing functions of the eigenvalues) as compared to low-pass filters implemented on the graph Laplacian, since we express the filters in terms of the adjacency matrix. The filter represented by the graph adjacency matrix corresponds to the identity (*y* = *x*) line.

As an alternate strategy to truncation (for RWR) or Taylor approximation (for HK), we here use polynomial interpolations of the filter functions [7]. While the ill-conditioned nature of the linear system to be solved for polynomial interpolation poses numerical challenges, we overcome these challenges using Arnoldi orthonormalization [6]. Polynomial interpolation improves the fidelity of the implemented polynomial filters to the filter function. Since polynomial interpolation is performed separately from the graph learning process, and the interpolation is agnostic to the input graph, the computational cost of computing the polynomial interpolation is negligible.

Our results on state-of-the-art scProteomic datasets demonstrate that scProfiterole enhances clustering performance and cell type annotation, outperforming established clustering algorithms in the clustering of single-cell transcriptomic data (K-means and Louvain [30]), as well as traditional GCN and contrastive learning baselines. We observe that the clustering performance of the spectral encoder, as well as the values of the learnable polynomial coefficients depend strongly on the spectral filter that is used for the initialization of these parameters. This observation suggests that, if the correct spectral filter is used to initialize the polynomial coefficients, this can significantly improve the robustness of the learning algorithms to fluctuations in the input data. Heat kernels clearly deliver the best performance in general, and polynomial interpolation via Arnoldi orthonormalization enhances the performance of Heat kernels over Taylor approximation. Altogether, scProfiterole provides a principled and scalable foundation for graph contrastive learning on noisy, high-dimensional single-cell proteomic data, spearheading the application of graph representative learning to extend the boundaries of next-generation proteomics.

## 2 Methods

### 2.1 Background and Problem Formulation

Len ***X*** = [ **x**_1_, …, **x**_*N*_ ] ∈ ℝ^*N* ×*d*^ denote the cell-to-protein abundance matrix, where ***X*** *i, j* represents the expression of protein *j* in cell *i, N* is the number of cells, and *d* is the number of proteins. As in scRNAseq analysis, the first step in the analysis of scProteomics data is to use ***X*** to construct a cell-to-cell similarity graph [23]. Here, since our focus is on the utilization of the cell-to-cell similarity graph for cell clustering and cell-type annotation, we assume that the cell-to-cell similarity graph is readily available as an undirected, unweighted graph 𝒢 ( 𝒱, ℰ) .Here, 𝒱 = { *v*_1_, *v*_2_, …, *v* _*N*_ } denotes the set of the cells and ℰ ⊆ 𝒱 × 𝒱, where ( *v*_*i*_, *v* _*j*_ ) ∈ ℰ indicates that cells *i* and *j* are similar to each other in terms of their protein expression profiles.

Li et al’s scPROTEIN framework [23] uses unsupervised deep graph contrastive learning (GCL) to learn cell embeddings from 𝒢 and ***X***. It comprises four key modules: 1) Data augmentation, 2) GCN-based encoder, 3) Node-level contrastive learning, 4) Alternating topology–attribute denoising. While effective, the standard GCN encoder primarily captures local neighborhood information and struggles with deeper architectures. As demonstrated in Fig. 1, addition of more than two neural network layers in the GCN encoder causes over-smoothing [5,22,35], leading to degradation of performance in downstream tasks. In addition, when the adjacency matrix is used for convolution, each *k*-hop neighborhood contributes equally to the propagation of features, making the framework vulnerable to the imperfections in the underlying graph. To overcome these issues, we decouple feature embedding from propagation using a Spectral GCN formulation, effectively replacing the adjacency matrix with a filtered version of the adjacency matrix in each layer.

### 2.2 Spectral Graph Convolution

Let *Â* = ***D***^−1/2^ ( ***A*** + ***I*** ) ***D***^−1/2^ denote the symmetrically normalized adjacency matrix of the cell-to-cell similarity graph, and ***X*** denote the cell-to-protein abundance matrix used as the node features input to the graph convolutional network. A Spectral GCN encoder [3,7,12,16] propagates the features using a spectral filter *g* ( ***Â*** ) in each layer *l* of the neural network:

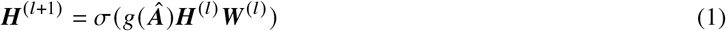

where *σ* (·) is an activation function, e.g., ReLU, ***W*** ^(*l*)^ is a learnable weight matrix, and ***H***^(0)^ = ***X***.

In practice, direct computation of *g* (***Â*** ) can be costly for arbitrary *g* (. ). The filter can be defined and evaluated on the eigenspectra of the graph adjacency matrix (Fig 2) or the graph Laplacian, but this is also costly since the computation of the eigen-decomposition of very large graphs is cost-prohibitive. Consequently, the filter is usually represented as a polynomial function:

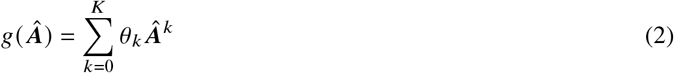

Here, *K* denotes the depth (number of hops) of the propagation on the graph and *θ*_*k*_s denote the polynomial coefficients used to adjust the relative weight of paths of different length in the graph. A significant benefit to the polynomial representation of filters is the ability to compute *g* (***Â*** ) ***H***^(*l*)^ as a series of sparse matrix-vector multiplications, without requiring explicit computation of *g* (***Â*** ), which usually is a dense matrix. Furthermore, setting *θ*_*k*_s as learnable parameters, the GCN can adaptively learn a filter that fits to the training data. Since the initialization of these parameters is essential, an explicit filter function is still required to initialize these parameters and guide the GCN. For initialization, earlier spectral GCNs use fixed coefficients such as random walks (*θ*_*k*_ = *α*^*k*^) [12,3]. Later approaches utilize different polynomial bases (e.g., Bernstein [16] or Chebyshev [15] polynomials) without explicit filter alignment. In recent work [6,7], we use polynomial interpolation to explicitly connect a spectral filter to a set of polynomial coefficients, enabling the use of any filter function.

### 2.3 Polynomial Interpolation of Filter Functions

The low-pass filters employed in this study are defined on the normalized adjacency matrix, whose eigenvalues lie within the range [− 1, 1 ]. These filters emphasize the low-frequency components of the graph spectrum, promoting smooth variations of node features across edges. This property aligns with the homophily assumption underlying clustering, where nodes within the same community are expected to share similar attributes.

Given a spectral filter *g* (*ω* ): [−1, 1 ] → ℝ and a desired polynomial degree *K*, we approximate *g* (*ω* ) with a polynomial expansion

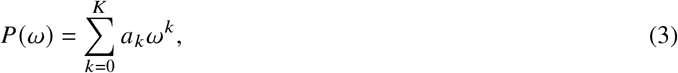

where the coefficients *a*_*k*_ serve as an approximation of the spectral response of the filter. These coefficients are subsequently used to initialize the parameters *θ*_*k*_ in the Spectral GCN encoder (Eq. 2).

To obtain stable and accurate approximations, we sample *K* spectral points in [−1, 1] using **Chebyshev nodes**:

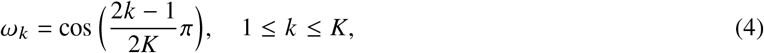

We evaluate the filter function *g* (·). at these Chebyshev nodes to obtain samples *g* (*ω*_*k*_ ) for 1 ≤ *k* ≤ *K*.

#### Vandermonde System

Given the samples, the coefficients of the interpolating polynomial are computed by solving

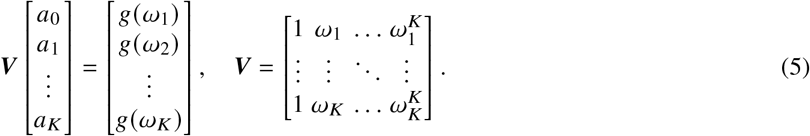

##### Algorithm 1

Coefficient Computation via Arnoldi Orthonormalization on **Ω**

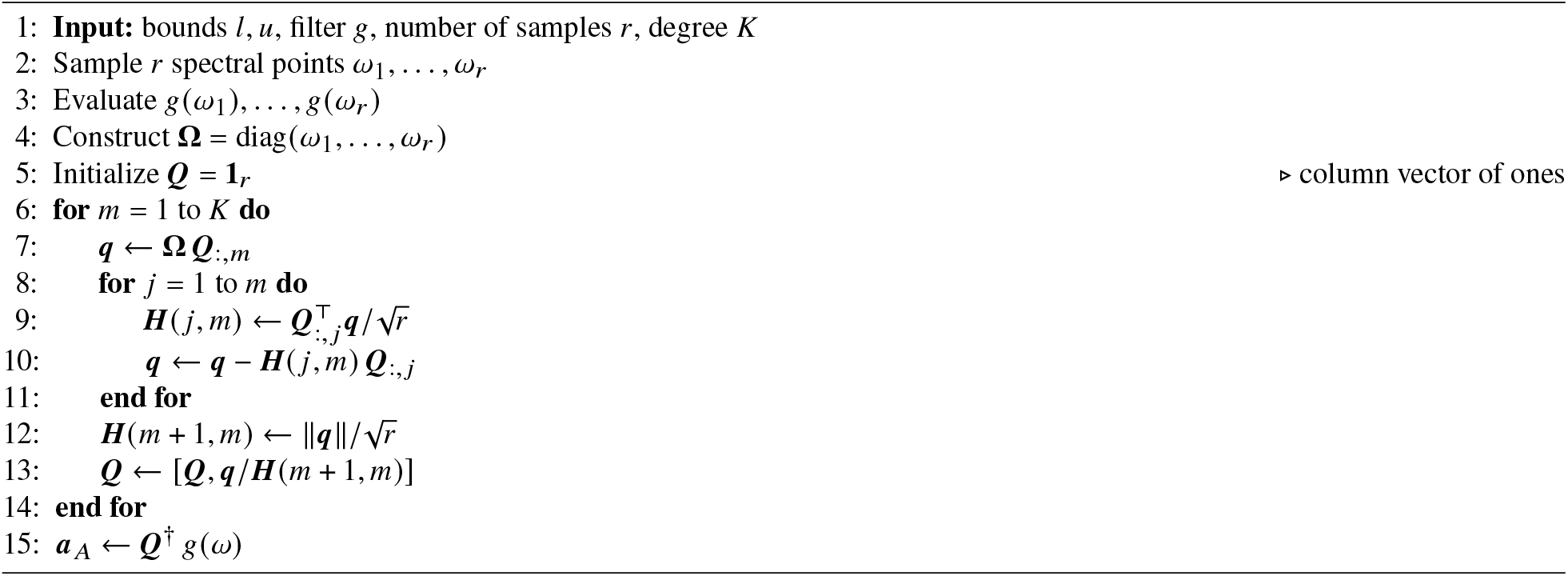

Since the Vandermonde matrix ***V*** is highly ill-conditioned [27], directly solving linear systems involving ***V*** can be numerically unstable. Even performing a QR decomposition, ***V*** = ***Q***_*V*_ ***R***_*V*_, does not guarantee stability:

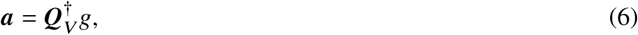

where 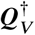 is the Moore–Penrose pseudoinverse. The fundamental reason for this instability is that Vandermonde matrices are closely related to Krylov subspaces, and when the nodes (or the corresponding eigenvalues) are clustered, the resulting Krylov subspace has nearly linearly dependent columns. As a result, standard QR decomposition algorithms fail to maintain numerical stability in practice [7].

#### Arnoldi-Orthonormalization

To overcome the aforementioned numerical stability issue, we apply QR decomposition to the diagonalized spectrum **Ω** = diag(*ω*_1_, …, *ω*_*K*_ ) using Arnoldi orthonormalization. This constructs an orthonormal basis ***Q*** _*A*_ for the Krylov subspace 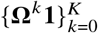, yielding:

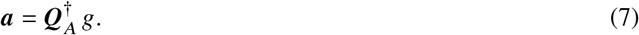

Note that, since the *ω*_*k*_s are real, Arnoldi orthonormalization here can be implemented using Lanczos’ algorithm. Specifically, we compute orthonormal matrix ***Q*** _*A*_, tridiagonal ***T***, and an almost-zero matrix ***Õ*** to satisfy:

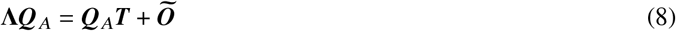

where **Λ** ∈ ℝ^*K*×*K*^, ***Q*** _*A*_ ∈ ℝ^(*K*+1)×*K*^, ***T*** ∈ ℝ^(*K*+1)×(*K*+1)^, and ***Õ*** ∈ ℝ^(*K*+1)×*K*^ is all zero except its last column. Then, as shown in [7], we can obtain a QR-decomposition for the Vandermonde matrix ***V*** as ***V*** ^(∗)^ = ***Q R***, such that ***V*** ^(∗)^ = ***V*** /∥ ***e*** ∥ and ***R*** = [ ***e***_1_, ***T e***_1_,, ·, ·, ·, ***T***^*K*^ ***e***_1_ ], where ***e***_1_ denotes the *K* + 1-dimensional vector of ones. Consequently, Eq. (7) provides a correct solution for the Vandermonde system of Eq. (5) through Eq. (6). Furthermore, it can be shown that the condition number of ***Q*** _***A***_ satisfies 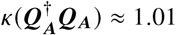. [6]. Therefore, Arnoldi orthonormalization, as detailed in Algorithm 1, yields accurate and numerically stable polynomial coefficients for interpolating the filter function.

### 2.4 Spectral Filters for Clustering Single-Cell Proteomics Data

In this section, we introduce three families of homophilic filters that are implemented in scProfiterole.

#### 1. Random Walk with Restart (RWR) Filter

The RWR filter is defined as:

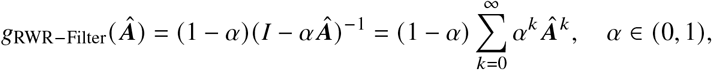

##### Algorithm 2

scProfiterole Convolution Layer (Encoder)

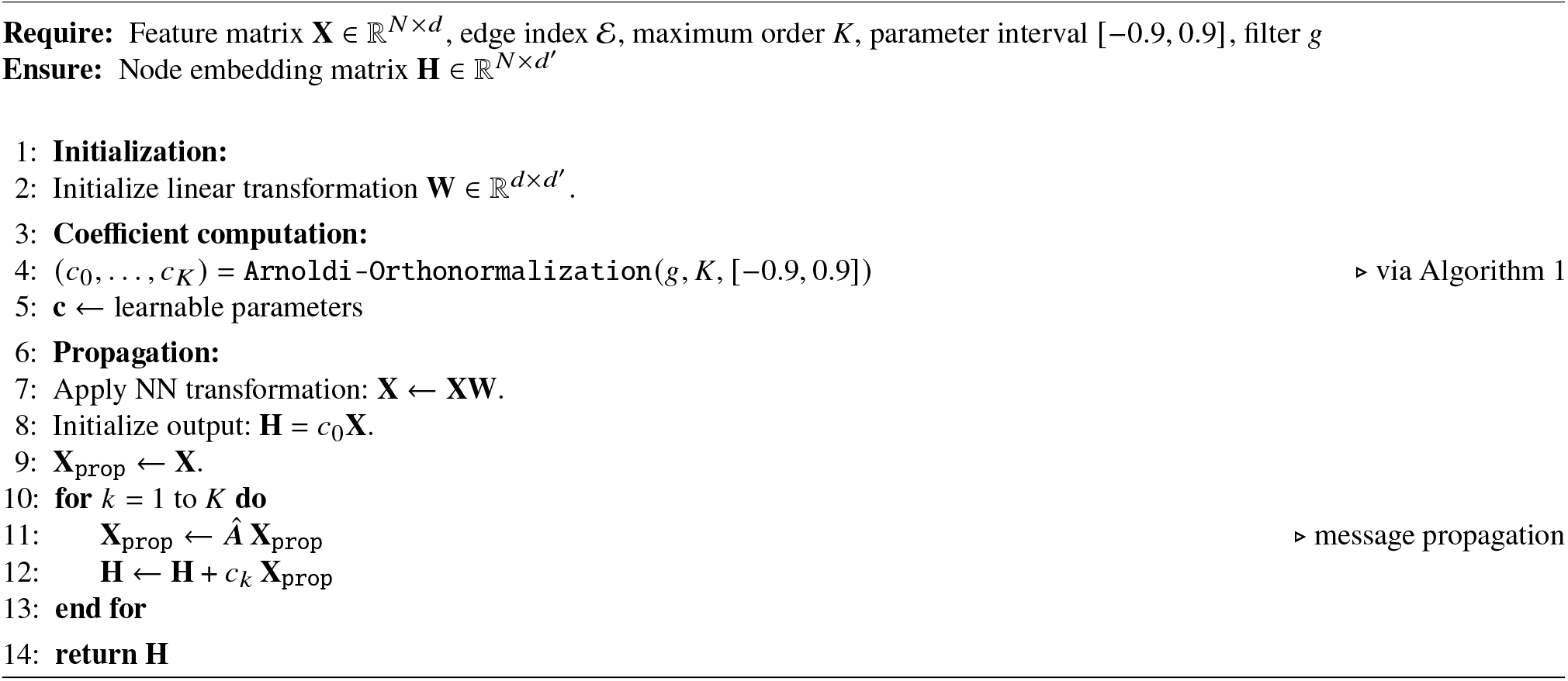

which is an infinite sum of the powers of the adjacency matrix, representing repeated neighborhood propagation with diffusion parameter *α*. While using RWR in GCNs, the infinite sum is truncated as:

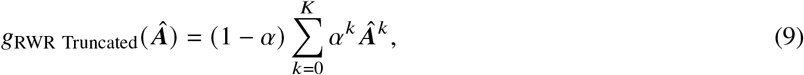

which corresponds to the *K*th iterate in the computation of the steady state matrix via power iteration. Therefore, the truncated sum may not provide a good approximation to *g*_RWR Filter_ ( ***Â***) for small values of *K*, which is often the case in GCN applications.

In contrast, the interpolated version of RWR we propose here uses the *K* + 1 polynomial coefficients (Eq. 2) that interpolate (Eq. 3) the following filter function, as computed by Arnoldi orthonormalization:

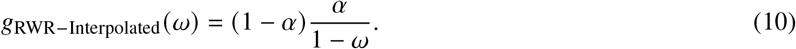

#### 2. Heat Kernel Filter

While the RWR filter is simple and interpretable, it is concentrated around a narrow band near the greatest eigenvalues of ***Â*** (Fig. 2, left). Different values of the parameter *α* provide little variation within this narrow band, limiting the flexibility of the RWR-filter in dealing with the sparse and noisy nature of cell-to-cell similarity graphs in scProteomics. Motivated by this insight, we also implement heat kernels in scProfiterole.

Let ℒ = ***I*** − ***Â*** denote the graph Laplacian. The heat kernel [21] is defined as:

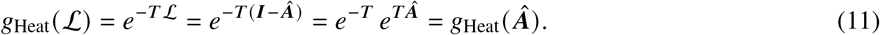

This can be thought of as a continuous random walk on the graph, where the parameter *T* specifies the duration of the walk. Kloster and Gleich [21] approximate the heat kernel using Taylor series:

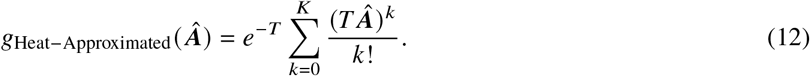

Since Taylor series approximates the function around a fixed point, it may deviate from the function that is being approximated for graphs whose eigenspectra are not concentrated around the fixed point. To this end, numerical polynomial interpolation may provide a more accurate approximation to the heat kernel. As above, we define the interpolated version of Heat kernel as the polynomial that uses the *K* + 1 polynomial coefficients (Eq. 2) that interpolate (Eq. 3) the following filter function, as computed by Arnoldi orthonormalization:

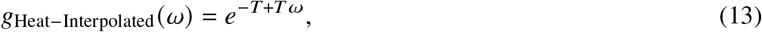

#### 3. Beta Kernel Filter

Beta distribution is commonly used as a wavelet basis for computer vision applications, but it also demonstrated promise as a kernel for anomaly detection on graphs [36]. While the general form of beta kernels can be used to define any type of filter on graph spectra, the zeroth-order Beta kernel provides a low-pass filter that can be directly represented as a polynomial of order *K*, where *K* serves a parameter (Fig 2, center). For this reason, zeroth order Beta kernel offers a desirable alternative for low-pass filter design, since it does not require truncation, approximation, or interpolation.

For a given *K*, the zeroth order Beta kernel is defined as follows:

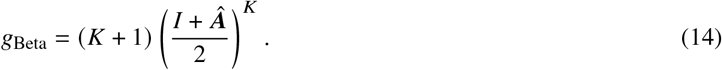

Observe that coefficients that correspond to this polynomial are the binomial coefficients normalized by 2^*K*^ */ K* **+** 1 . We implement the beta kernel by directly incorporating these coefficients as the initial values of the coefficients in the spectral filter (Eq. 2). The proposed encoder details is given in Algorithm 2, full mathematical procedure can be found in Appendix subsection **??**.

In summary, we introduce a polynomial interpolation based framework to implement spectral graph filters, enabling scalable and stable low-pass filtering of node features in GCN-Encoders. Filter coefficients are computed using Chebyshev nodes and stabilized via Arnoldi orthonormalization, overcoming the ill-conditioning of standard Vandermonde systems. We implement three homophilic filters: (i) Random Walk with Restart (RWR) for repeated neighborhood propagation, (ii) Heat Kernel for continuous graph diffusion, and (iii) zeroth-order Beta Kernel as a direct low-pass polynomial. For RWR and Heat kernels, polynomial interpolation ensures accurate and stable approximation, while the Beta kernel provides a straightforward low-pass design, collectively enhancing node feature smoothness and cluster coherence in single-cell proteomics data.

## 3 Results

In this section, we first detail the datasets and experimental setup. We then present the performance of the spectral graph filters implemented by scProfiterole in the context of clustering scProteomics data and cell type annotation. Subsequently, we investigate the differences between the spectral filters in terms of how they initialize the polynomial coefficients, how initialization influences the learned coefficients, and what filter functions correspond to the learned coefficients. We then discuss the insights derived from these results on the design of spectral filters that are best suited to the clustering of cell-to-cell similarity graphs derived from scProteomics data.

### 3.1 Datasets and Experimental Setup

We evaluate scProfiterole in the context of clustering cell-to-cell similarity graphs derived from scProteomics data and cell type annotation.

#### Datasets and Preprocessing of scProteomics Data

The datasets we use for this purpose are shown in Table 1. To comprehensively compare the performance of the spectral filters implemented in scProfiterole and investigate the effect of parameters, we use the `Scope2_Specht` dataset [33], profiling 3,042 proteins across 1,490 cells. To test robustness against batch effects and assess the generalizibility of our conclusions to other datasets, we also obtain a second dataset by integrating two relatively smal datasets, N2 [9] (108 cells, 1,068 proteins) and nanoPOTS [9] (61 cells, 1,225 proteins). We use the 762 overlapping proteins between these two datasets to construct a shared protein expression data matrix, and construct the cell-to-cell similarity graph using this integrated data matrix. To obtain the cell-to-cell similarity graph, we utilize the framework introduced by scPROTEIN [23], performing peptide-level uncertainty estimation via multitask heteroscedastic regression and uncertainty-weighted aggregation to obtain noise-reduced protein abundances, . followed by the computation of the pairwise correlations between the protein expression profiles of the cells.

**Table 1:** scProteomics datasets used in our experiments.

| Name | # Cells | # Proteins | Cell type (#) |
| --- | --- | --- | --- |
| Scope2_Specht | 1490 | 3042 | Monocytes (394) and Macrophages (1096) |
| N2 | 108 | 1068 | C10 (36), RAW (36), SVEC (36) |
| nanoPOTS | 61 | 1225 | C10(19), RAW(22), SVEC(20) |

#### Performance Criteria

To assess the performance of the filters in clustering and cell-type annotation, we use four standard performance criteria: Adjusted Rand Index (ARI) [17], Average Silhouette Width (ASW) [31], Normalized Mutual Information (NMI) [11], and Purity Score (PS) [26]. The definitions of these measures are provided in Appendix **??**. For all of these measures, higher values indicate more desirable performance. We repeat each experiment eight times using different random seeds and report the Mean ± 95% Confidence Interval for all performance figures. **Algorithms and Baselines**. scProfiterole implements three spectral GCN based encoders for GCL-based clustering of scProteomics data: (i) Random Walk with Restart (**RWR**) filters, (ii) **Heat** kernel filters, and (iii) **Beta kernel** filters. For the RWR filter, we consider the **truncated** (standard) and **interpolated** (proposed) versions. For the Heat kernel, we consider the **approximated** (standard) vs. **interpolated** (proposed) versions. Since Beta Kernel naturally employs a finite polynomial basis, we consider its **direct** implementation. We compare the performance of these three algorithms against three types of baselines: 1) As classical algorithms for clustering single cell omic data, **K-means** clustering and **Louvain** [30] algorithm for community detection. For both algorithms, we apply dimensionality reduction via Principal Component Analysis (**PCA**). 2) As a baseline for **GCN**-based encoding, GCL-based clustering algorithm that uses the adjacency matrix of the cell-to-cell similarity graph for graph encoding [23]. 3) As a baseline for spectral encoding, scProfiterole with **random initialization** of polynomial coefficients (without filter guidance).

#### Parameter Settings

The families of spectral filters we implement have one parameter in common: The degree *K* of the polynomial that is used to implement the filter. While this is the sole parameter for the Beta kernels and random initialization, Random walk filters and Heat kernels each have one additional parameter: Restart probability (*α*) for Random walk, diffusion time (*t*) for Heat kernels. In our first round of experiments we fix *K* = 6 for the Random walk and Heat kernel filters, while we vary *K* from 2 to 6 for the Beta kernels. We choose this value of *K* as we expect the cell-to-cell similarity graphs to be small world networks (thus a smaller *K* can be desirable) and we would like to maximize the expressive power of the polynomial approximation/interpolation (thus a larger *K* can be desirable). Regardless, we perform additional experiments for values of *K* up to 24 to fully characterize the effect of the polynomial degree on the performance of spectral filters. For random walks, we use five equispaced values of *α* in the range [ 0.05, 0.95 ]. For the heat kernel, we consider integer values of diffusion time *T* ranging from 1 to 5. Finally, for the primary experiments reported here, we set the number of dimensions for cell embeddings to *D* = 64, but we also investigate the effect of *D* for values ranging from 32 to 256. Upstream, for the construction of the cell-to-cell similarity matrix, we use the algorithm implemented by scProtein [23]. This algorithm uses the parameter *h* as the threshold on correlation for an edge to be included between two cells. In the first set of experiments reported here, we set *h* = 0.15. This leads to a graph with 274, 816 edges for the `Scope2_Specht` dataset and a graph with 4, 950 for the integrated `N2` and `nanoPOTS` dataset. At the end of this section, we assess the effect of *h* (i.e., the sparsity of the cell-to-cell similarity matrix) on clustering performance.

### 3.5 Performance Evaluation

We first compare the clustering performance of the Random Walk with Restart (RWR) filters, Heat Kernel filters, and Beta Kernel filters against baselines on the `Scope2_Specht` dataset. The results of this analysis are shown in Figure 3. **GCN encoders and low-pass filters outperform classical algorithms and random initialization**. We observe that classical baselines (K-means and Louvain) consistently underperform GCL-based algorithms. In particular, PCA+Louvain exhibits the weakest performance. GCL using adjacency matrix for GCN-based encoding outperforms classical algorithms with respect to all performance criteria, suggesting that graph contrastive learning adds value to the clustering of scProteomics data. We observe that spectral encoders with random initialization of polynomial coefficients deliver improved performance with increasing *K* (degree of the polynomial representing the filter), also outperforming classical algorithms. However, their performance remains below that of the GCN encoder that uses adjacency matrix according to all criteria, and well below all spectral encoders that use RWR, heat kernel, or beta kernel filters for initialization. This observation highlights that the design of appropriate filters is essential for spectral encoders to improve performance over standard GCNs.

**Fig. 3:**
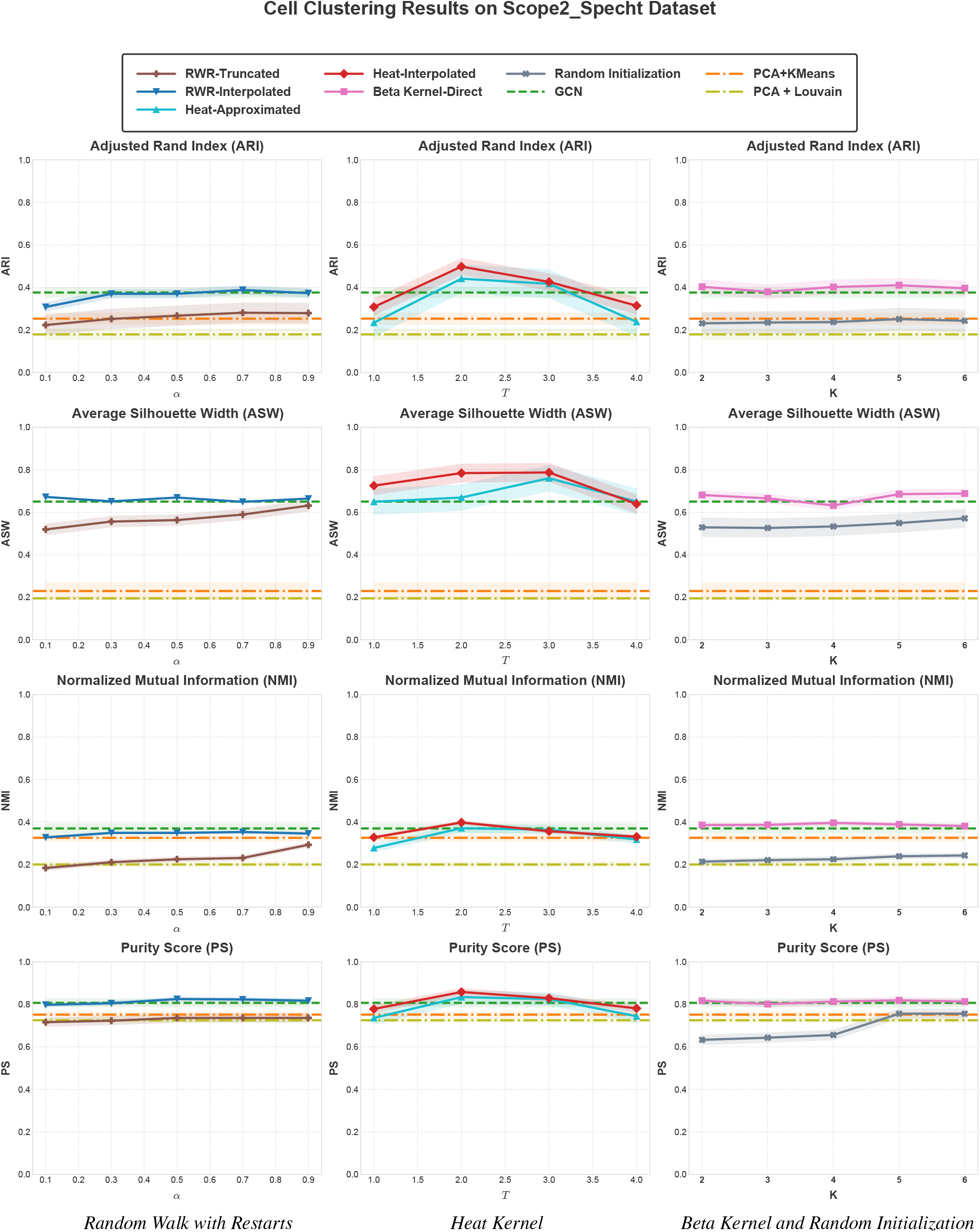
Clustering performance of the spectral filters implemented in scProfiterole on the Scope2_Specht dataset. Four performance criteria are shown as a function of restart probability (*α*) for random-walk based filters (Left), diffusion time (*T* ) for Heat kernel filters (Middle), and polynomial degree (*K*) for the Beta kernel and Random Initialization (Right). For random walk and heat kernel, *K* = 6. Plot shows mean ± 95% confidence interval over eight different runs with random seeds. The performance of other baselines (GCL with GCN encoder that uses the adjacency matrix for convolution, PCA+K-means, PCA+Louvain) does not depend on a parameter and they are shown on all panels to facilitate comparison.

#### Heat kernels deliver the best clustering performance

As seen in Figure 3, the Heat kernel based spectral filters outperform other filters, delivering optimal performance for *T* ∈ [ 2, 3 ]. For *T* = 2, as compared to the baseline GCN that uses the adjacency matrix for convolution, Interpolated Heat kernel provides 29.7% improvement in ARI, 9.1% improvement in ASI, 8.1% improvement in NMI, and 3.7% improvement in PS. RWR filter delivers inferior performance as compared to the GCN baseline, but its performance is improved with increasing polynomial degree (*K*), as would be expected. Interpolation makes RWR competitive with the baseline and the effect of *K* vanishes with interpolation. Similarly, the Beta kernel delivers performance that is competitive with the baseline, with little dependence on *K*. These results show that the performance of spectral graph filters in clustering scProteomics data depends highly on the choice of the spectral filter. While interpolated Heat kernels provide consistent and considerable improvement over the GCN encoder that uses the adjacency matrix, other filters either fail to deliver meaningful improvement or have a degrading effect on clustering performance.

#### Interpolation via Arnoldi orthonormalization enhances spectral filter performance

For both the RWR and Heat kernel filters, we observe considerable performance gap between the interpolated vs. truncated (for RWR) or approximated (Heat kernel) filters. While the performance of the RWR filter is improved with increasing order of the polynomial representation of the filter (*K*), interpolation removes this dependency. This observation suggests that interpolation of spectral filters enables working with polynomials of smaller order, capturing higher order relationships in the graph by considering shorter paths than those needed by truncated polynomials. Interpolated filters accomplish this by choosing the polynomial coefficients (weights assigned to path lengths) to approximate the true filter with fidelity. Interpolation also results in drastic improvement in the performance of the Heat kernel, making Heat kernels the best choice of spectral filters for clustering scProteomics data. For Heat kernel based filters, the dependency on diffusion time *T* is consistent between the approximated and interpolated versions, where clustering performance reaches its peak at *T* = 2 and declines for *T >* 3. These results suggest that the ability to interpolate spectral filters using a numerically stable framework enables spectral filters to realize their potential in the context of clustering scProteomics data.

#### The learned filter strictly depends on the initialization of the polynomial coefficients

In Fig. 4, the initial values of the polynomial coefficients, the learned values of the polynomial coefficients, and the filter function that corresponds to the learned polynomial are shown for the setting that results in the optimal performance for each filter family on the `Scope2_Specht` dataset. Comparison of the first two rows of this figure suggests that the learned polynomial coefficients heavily depend on their initial values. This result suggests that the optimization of the polynomial coefficients alongside neural network parameters may be hampering the search in the coefficient space. To this end, at the absence of dedicated learning algorithms that optimize polynomial coefficients more effectively, the choice of the spectral filter for initialization remains a significant hyperparameter. This conclusion is reinforced by the filters that result from random initialization (Fig. 4, right panel). Different random initializations lead to noticeably different learned polynomial coefficients and, consequently, to substantially different learned filter shapes, despite using the same model and training protocol. Taken together, these observations suggest that the initialization of coefficients with explicit filters guides the learning of polynomial coefficients toward a desired filter function and makes the learning of these parameters less sensitive to specific training data.

**Fig. 4:**
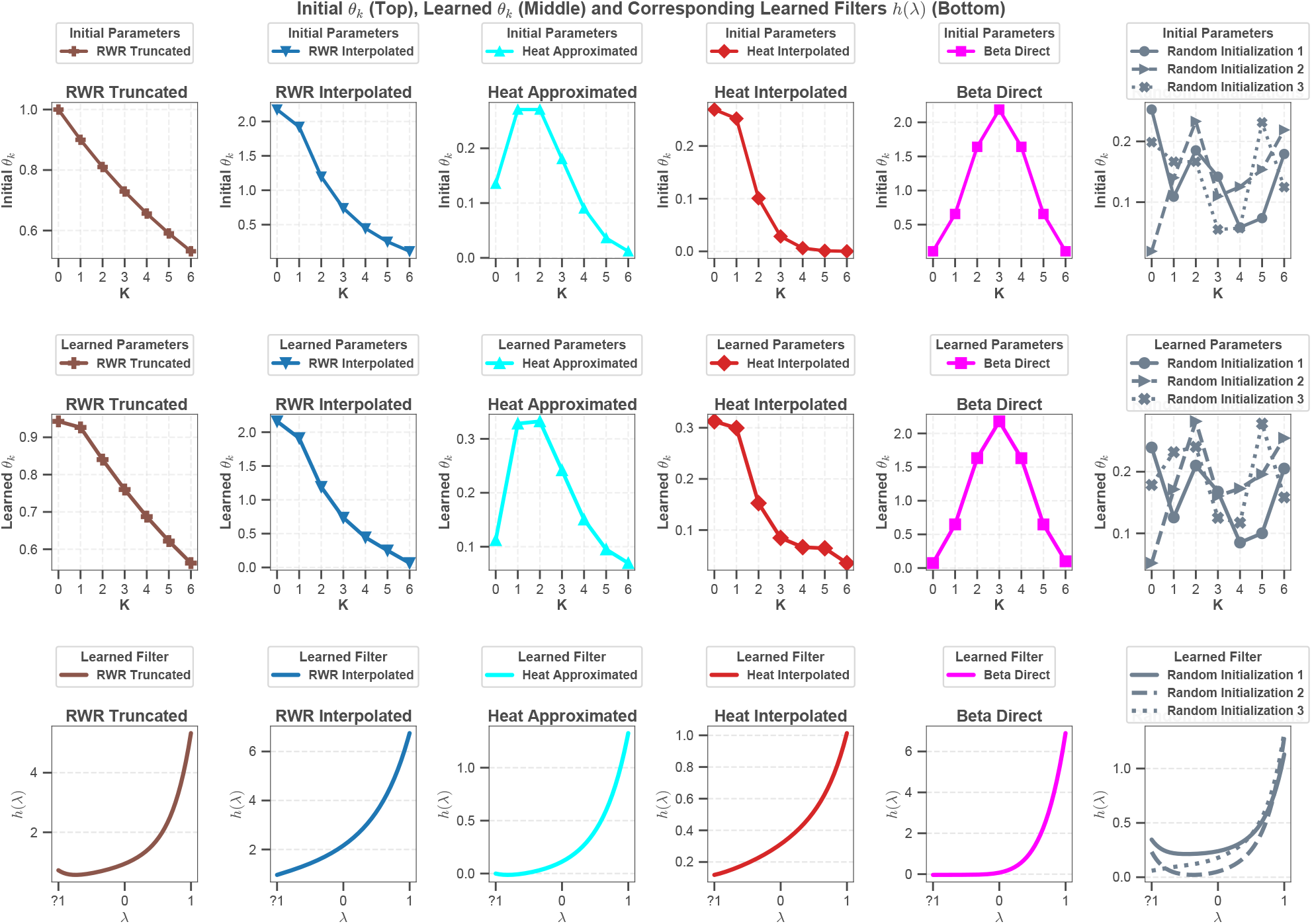
Learned spectral coefficients and corresponding filters on the Scope2_Specht dataset. Comparison of initial and learned polynomial coefficients *θ*_*k*_ (top and middle rows, respectively) and filter function *g* (*λ* ) corresponding to the learned coefficients (bottom row) across five spectral filters implemented in scProfiterole: Random Walk with Restart (Truncated and Interpolated), (Direct), Heat Kernel (Approximated and Interpolated), and Beta Kernel, as well as three samples of randomly initialized coefficients. In these experiments, *K* = 6 for all filters, *α* = 0.9 for the RWR filter, and *T* = 2 for Heat kernel.

#### Low-pass filters that are effective in clustering scProteomics data provide a broader view of the eigenspectra

The last row of Fig. 4 shows the filter function that corresponds to the learned polynomial coefficients for each filter family. While all functions represent low-pass filters (recall that these functions are defined on the eigenvalues of the normalized adjacency matrix), the curves for the interpolated Heat kernel and Random walk filters represent a unique pattern. Unlike other filters that suppress negative eigenvalues, these filters permit negative eigenvalues, with preference to eigenvalues with smaller magnitude. For positive eigenvalues of the adjacency matrix, these filters assign more importance to eigenvalues with larger magnitude, but the rise of the curve is not as sharp or pronounced as other filters. Inspecting the initial and learned polynomial coefficients in the first and second rows of the figure, we observe that these filters accomplish this by assigning more weight to the zeroth and first order terms, while the weights of higher order terms decline superlinearly for these filters.

#### Heat kernels effectively trade-off sparsity and noise

To investigate the effect of the sparsity-noise trade-off on the performance of GCL algorithms for clustering scProteomics data, we perform experiments by varying the sparsity of the cell-to-cell similarity matrix. For this purpose, we adjust the parameter *h* of ScProtein [23], which is the threshold on the correlation between two cell embeddings to retain an edge between two cells. The results of this experiment are shown in Fig. 5. to assess the effect of the sparsity of the network on performance by adjusting the threshold on retaining edges in the cell-to-cell similarity graph. In general, we observe that the differences between filters in clustering are mostly preserved across different levels of sparsity. The optimal level of sparsity appears to be consistent for best-performing algorithms; GCN with adjacency matrix and Heat kernels deliver optimal performance for *h* ∈ [ 0.15, 0.3 ]. For these algorithms, increasing graph density further degrades performance. On the other hand, clustering performance degrades drastically with increasing sparsity across the board. Nevertheless, Heat kernels appear to be most robust against increasing sparsity, and approximated Heat kernels outperform interpolated Heat kernels on sparser graphs. This may be because of Taylor approximation’s improved fidelity to the heat kernel on sparser graphs, since the eigenspectra of the adjacency matrix tends to get clustered around the origin with increasing sparsity [10].

**Fig. 5:**
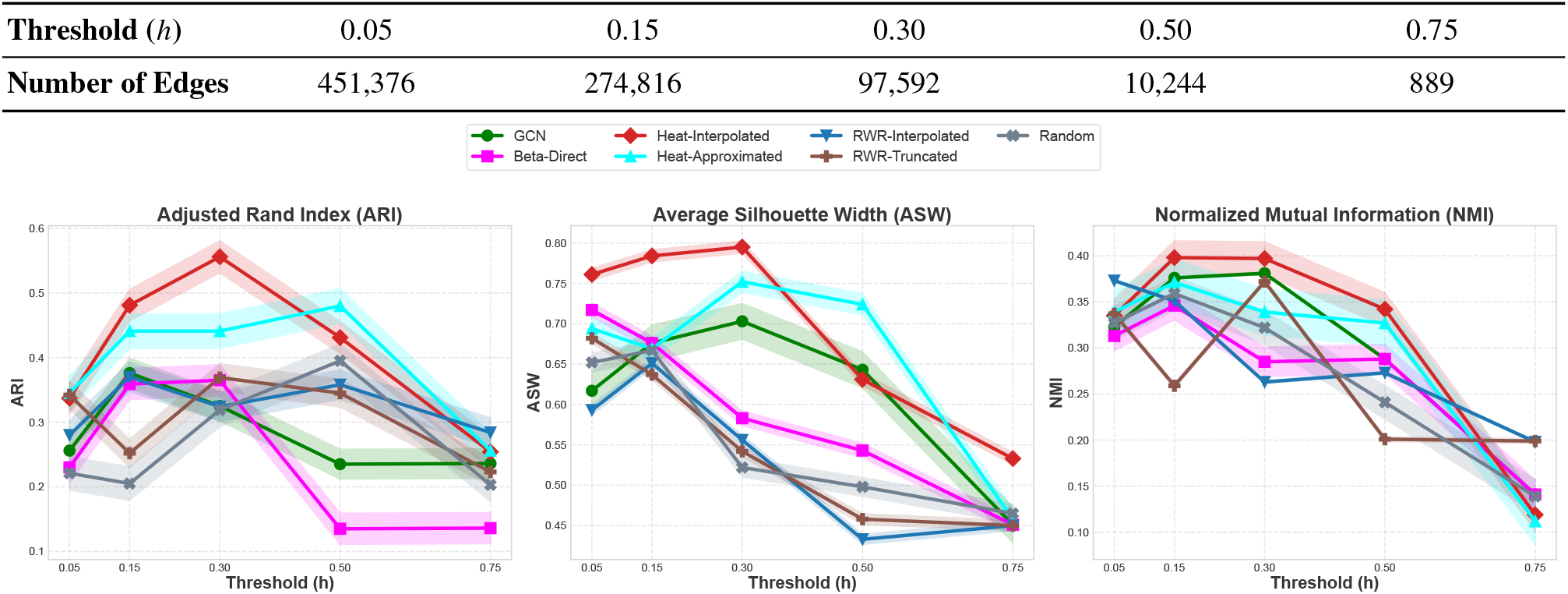
The effect of the sparsity of the cell-to-cell similarity graph on the clustering performance of GCL algorithms. Clustering performance of each algorithm is shown for performance criteria ARI, ASW, and NMI, as a function of the threshold on the correlation of cell embeddings *h* ∈ { 0.05, 0.15, 0.3, 0.5, 0.75 } on the `Scope2_Specht` dataset. For spectral filters, polynomial degree is set to *K* = 6 in these experiments. The number of edges in the cell-to-cell similarity graph is given in the top panel. The graph gets sparser with increasing *h*. For all performance criteria, mean ± 95% confidence interval over eight different seed runs are shown.

#### Spectral filters and interpolation incur negligible computational cost within the GCL framework

To assess the computational burden introduced by the spectral encoder and polynomial interpolation of spectral filters, we comprehensively assess the runtime of all algorithms on the Scope2_Specht. These experiments are performed in the Google Colab environment, with NVIDIA A100-SXM4 GPU with 40 GB of HBM2 VRAM, complemented by 83.5 GB of system RAM and 235.7 GB of available disk storage. We observe that classical algorithms (K-means and Louvain) are much fater than GCL algorithms, performing PCA and clustering within two seconds. In contrast, GCL algorithms spend close to an hour to cluster the `Scope2_Specht`. A significant fraction of this time is devoted to Phase 1 training (before graph construction) and Phase 2 training (after cell embeddings are computed), whereas the training of the graph encoder takes less than a second for the GCN encoder that uses the adjacency matrix, as well as the spectral encoders. The time required to compute the polynomial interpolation is also in the order of miliseconds, demonstrating that the use of spectral encoders and polynomial interpolation do not add any additional burden to the GCL. Detailed results comparing the runtime of different algorithms are shown in Table **??** of Supplementary Materials.

#### The key observations are replicated on a second dataset

To assess the generalization of our observations and the robustness of the spectral filters to heterogeneous data sources, we also evaluate the clustering performance of spectral filters on the integrated N2 and nanoPOTS datasets. Fig. 6 shows the Average Silhouette Width (ASW) of the clustering delivered by each filter family as a function of its key parameters. Since this dataset is smaller than the `Scope2_Specht`, the performance of the algorithms exhibit greater variance on this dataset. As observed on the `Scope2_Specht` dataset, GCL-based algorithms outperform classical algorithms. Also consistent with the `Scope2_Specht` dataset, we observe that the interpolated Heat kernel delivers best clustering performance, with peak performance for *T* ∈ [ 2, 3 ]. We also observe that the interpolated filters outperform truncated (for Random walk) and approximated (for Heat kernel) versions of the filters. The cell clustering provided by the truncated Random walk filter is of lower quality than that obtained using the GCN encoder with the adjacency matrix, but the quality of clustering improves with increasing order of the polynomial filter. Arnoldi orthonormalization based interpolation helps the Random walk filter deliver better clustering than the baseline, while also removing the dependency on *K*. The cell clustering provided by the Beta kernel is competitive with the baseline, with no clear dependency on *K*. Random initialization exhibits higher variance and becomes competitive for greater polynomial degree (*K*), suggesting that spectral encoders can learn polynomial coefficients more effectively on smaller datasets, provided that the polynomial has a sufficiently high degree.

**Fig. 6:**
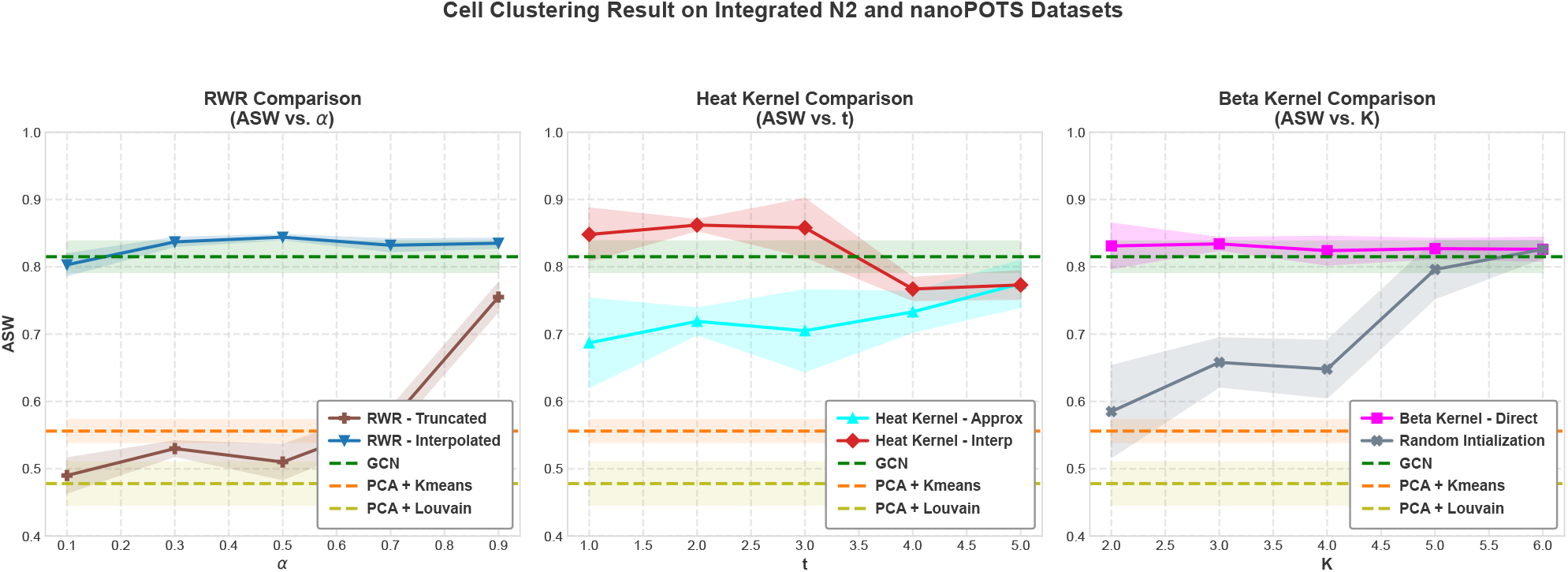
Cell clustering performance of the spectral filters on the integrated N2 and nanoPOTS datasets. ASW scores are reported for Truncated vs. Interpolated Random walk filters as a function of restart probability *α* (left), Beta kernel as a function of the order *K* of the polynomial filter (center), and Approximated vs. Interpolated Heat kernel as a function of diffusion time *T* (right). The figure shows Mean ± 95% Confidence Interval over eight different seed runs.

## 4 Conclusion

We present scProfiterole, a spectral graph contrastive learning framework tailored to clustering single-cell proteomics data. By utilizing learnable low-pass spectral filters including random walk, heat, and beta kernels, and efficient Arnoldi orthonormalization-based polynomial interpolation, scProfiterole overcomes challenges of graph sparsity, noise, and oversmoothing without requiring deep network architectures. Experiments on state-of-the-art scProteomics datasets demonstrate that scProfiterole enhances clustering and cell type annotation, with heat kernel initialization providing the most consistent improvements. Overall, scProfiterole integrates the interpretability of spectral graph theory with the flexibility of contrastive learning, providing a scalable and principled foundation for robust representation learning in high-dimensional, noisy single-cell proteomic data.

## Supporting information

https://github.com/mustafaCoskunAgu/scProfiterole/blob/main/Single_Cell_Proteomics__RECOMB_2026_Supplementary.pdf

## Acknowledgments

This work was supported in part by US National Institutes of Health (NIH) grant R01-LM012980 from the National Library of Medicine. The authors would like to thank Kaan Yorgancıoğlu (CWRU), Kasey Wei (CWRU), Willis Erdman (CWRU), Aidan Leblanc (CWRU), Abdelkader Baggag (Qatar Computing Research Institute), and Ananth Grama (Purdue) for many useful discussions.

## Disclosure of Interests

The authors have no competing interests to declare that are relevant to the content of this article.

## Notes

https://github.com/mustafaCoskunAgu/scProfiterole

### Competing Interest Statement

The authors have declared no competing interest.

### Summary of Updates

Based on the rebutal in Recomb 2026, new figures and subsections are added. For example, Fig 5 has been added. All other figures are presented with 8 run 95 % confidence interval.

