## Supplementary material for "scProfiterole: Clustering of Single-Cell Proteomic Data Using Graph Contrastive Learning via Spectral Filters": https://github.com/mustafaCoskunAgu/scProfiterole/blob/main/Single_Cell_Proteomics__RECOMB_2026_Supplementary.pdf

**Abstract.** Novel technologies for the acquisition of protein expression data at the single cell level are emerging rapidly. Although there exists a substantial body of computational algorithms and tools for the analysis of single cell gene expression (scRNAseq) data, tools for even basic tasks such as clustering or cell type identification for single cell proteomic (scProteomics) data are relatively scarce. Adoption of algorithms that have been developed for scRNAseq into scProteomics is challenged by the larger number of drop-outs, missing data, and noise in single cell proteomic data. Graph contrastive learning (GCL) on cell-to-cell similarity graphs derived from single cell protein expression profiles show promise in cell type identification. However, missing edges and noise in the cell-to-cell similarity graph requires careful design of convolution matrices to overcome the imperfections in these graphs. Here, we introduce scProfiterole (Single Cell Proteomics Clustering via Spectral Filters), a computational framework to facilitate effective use of spectral graph filters in GCL-based clustering of single cell proteomic data. Since clustering assumes a homophilic network topology, we consider three types of homophilic filters: (i) random walks, (ii) heat kernels, (iii) beta kernels. Direct implementation of these filters is computationally prohibitive, thus the filters are either truncated or approximated in practice. To overcome this limitation, scProfiterole uses Arnoldi orthonormalization to implement polynomial interpolations of any given spectral graph filter. Our results on comprehensive single cell proteomic data show that (i) graph contrastive learning with learnable polynomial coefficients that are carefully initialized improves the effectiveness and robustness of cell type identification, (ii) heat kernels and beta kernels improve clustering performance over adjacency matrices or random walks, and (iii) polynomial interpolation of spectral filters outperforms approximation or truncation. The source code for scProfiterole and Supplementary Materials are available at `https://github.com/mustafaCoskunAgu/scProfiterole`.

**Keywords:** Single cell proteomics, Clustering, Cell type identification, Graph contrastive learning, Spectral graph filters, Polynomial Interpolation

### Appendix

#### scProfiterole: Clustering of Single-Cell Proteomic Data UsingGraph Contrastive Learning via Spectral Filters

Mustafa Coşkun, Filipa Blasco Lopes, Pınar Tolunay Kubilay, Mark R. Chance, Mehmet Koyutürk

#### A Mathematical Details of scProfiterole

##### A.1 Data augmentation.

Contrastive learning seeks to learn invariant representations by comparing similar and dissimilar data pairs. To construct similar (positive) pairs, we generate multiple augmented views of the input graph using two augmentation strategies: *edge dropping* [11] and *feature masking*. Although a variety of contrastive mechanisms exist—such as topology-aware graph views proposed in [4,10] in addition to edge dropping—we focus exclusively on *edge dropping* [11] and *feature masking* to maintain consistency with the scPROTEIN framework. Exploring these alternative augmentation techniques is left for future work.

**Edge dropping.** Given an edge-dropping ratio $p_{\text{de}}$, we randomly remove edges from $E$. Specifically, we sample an indicator matrix $R \in \mathbb{R}^{N \times N}$ such that

$$\boldsymbol{R}_{ij} \sim \begin{cases} \text{Bernoulli}(1 - p_{\text{de}}), & \boldsymbol{A}_{ij} = 1, \\ 0, & \boldsymbol{A}_{ij} = 0, \end{cases}$$

and obtain the perturbed adjacency matrix

$$\widetilde{\boldsymbol{A}} = \boldsymbol{A} \odot \boldsymbol{R},$$

where $\odot$ denotes the Hadamard product.

**Feature masking.** To mask node features, we sample a masking vector $\boldsymbol{M} \in \mathbb{R}^{N \times 1}$ with entries

$$\boldsymbol{M}_i \sim \text{Bernoulli}(1 - p_{\text{mf}}),$$

where $p_{\text{mf}}$ is the masking probability. The masked feature matrix is then given by

$$\widetilde{\boldsymbol{X}} = \begin{bmatrix} \mathbf{x_1} \odot \boldsymbol{M} \\ \mathbf{x_2} \odot \boldsymbol{M} \\ \vdots \\ \mathbf{x_N} \odot \boldsymbol{M} \end{bmatrix},$$

where $[\cdot;\cdot]$ denotes row-wise vector concatenation.

In each training iteration, these augmentation procedures produce two correlated graph views,

$$\mathcal{G}_1 = (\mathcal{V}, \widetilde{\mathcal{E}}_1, \widetilde{\boldsymbol{X}}_1), \qquad \mathcal{G}_2 = (\mathcal{V}, \widetilde{\mathcal{E}}_2, \widetilde{\boldsymbol{X}}_2),$$

which are used for contrastive learning.

##### A.2 Arnoldi-orthonormalization–based spectral graph encoder.

Given the augmented graph views $\mathcal{G}_1$ and $\mathcal{G}_2$, we learn cell embeddings via node-level graph contrastive learning using an Arnoldi-guided spectral encoder. The standard GCN [9] serves as the baseline encoder, where an $L$-layer GCN updates node representations through neighborhood aggregation:

$$\boldsymbol{Z}^{(l+1)} = \sigma\left(\hat{\boldsymbol{A}} \boldsymbol{Z}^{(l)} \boldsymbol{W}^{(l)}\right), \tag{1}$$

with $\hat{\boldsymbol{A}} = \boldsymbol{D}^{-1/2}(\boldsymbol{A} + \boldsymbol{I})\boldsymbol{D}^{-1/2}$ denoting the normalized adjacency, $\boldsymbol{W}^{(l)}$ the trainable weights, and $\sigma(\cdot)$ a (parametric) ReLU. However, as illustrated in Fig. **??**, deep GCNs suffer from over-smoothing, motivating the use of spectral propagation.

A spectral GCN encoder [1,3,5,8] applies a spectral filter $g(\hat{\boldsymbol{A}})$ at each layer:

$$\boldsymbol{H}^{(l+1)} = \sigma\Big(g(\hat{\boldsymbol{A}})\boldsymbol{H}^{(l)}\boldsymbol{W}^{(l)}\Big), \qquad \boldsymbol{H}^{(0)} = \boldsymbol{X}. \tag{2}$$

Direct computation of $g(\hat{\boldsymbol{A}})$ via eigendecomposition is impractical on large graphs; therefore the filter is implemented as a $K$-th order polynomial:

$$g(\hat{\boldsymbol{A}}) = \sum_{k=0}^{K} \theta_k \hat{\boldsymbol{A}}^k, \tag{3}$$

allowing efficient evaluation through iterative matrix–vector multiplications. The coefficients $\theta_k$ become learnable filter parameters. Earlier spectral GCNs used fixed choices (e.g., random-walk $\theta_k = \alpha^k$ [5,1]) or predefined bases (Bernstein [8], Chebyshev [7]). Our recent work [2,3] instead employs polynomial interpolation to explicitly align any desired spectral filter with its polynomial coefficients, enabling Arnoldi-guided spectral learning (Algorithm **??**). Note that the main text introduces three families of filters $g(\cdot)$: (i) `RWR-Truncated` and `RWR-Interpolated` (Eq. **??**, **??**), (ii) `Heat-Approximated` and `Heat-Interpolated` (Eq. **??**, **??**), and (iii) the `Beta-Direct` kernel (Eq. **??**).

For both augmented views, we use a weight-sharing Arnoldi-based spectral GNN (Algorithm **??**) to obtain embeddings $\boldsymbol{H}_1$ and $\boldsymbol{H}_2$, with all layer dimensionalities kept consistent across views.

##### A.3 Node-level graph contrastive learning.

The embeddings $\boldsymbol{H}_1$ and $\boldsymbol{H}_2$ from the two augmented views are passed through a shared projection head $\phi(\cdot)$ (a two-layer MLP) that maps them into a common latent space where the contrastive loss is computed. For each node $i$, its projected embeddings $\phi(\boldsymbol{h}_{1i})$ and $\phi(\boldsymbol{h}_{2i})$ form a positive pair, while all other node embeddings act as negatives.

We use cosine similarity $\cos(\cdot,\cdot)$ and the InfoNCE loss. For a positive pair $(\boldsymbol{h}_{1i}, \boldsymbol{h}_{2i})$, the node-level contrastive loss is

$$\ell(\boldsymbol{h}_{1i}, \boldsymbol{h}_{2i}) = -\log \frac{\exp(\theta(\boldsymbol{h}_{1i}, \boldsymbol{h}_{2i})/\tau)}{\exp(\theta(\boldsymbol{h}_{1i}, \boldsymbol{h}_{2i})/\tau) + \sum_{j\neq i}\big[\exp(\theta(\boldsymbol{h}_{1i}, \boldsymbol{h}_{1j})/\tau) + \exp(\theta(\boldsymbol{h}_{1i}, \boldsymbol{h}_{2j})/\tau)\big]}, \tag{4}$$

where $\theta(\boldsymbol{h}_{1i}, \boldsymbol{h}_{2i}) = \cos(g(\boldsymbol{h}_{1i}), g(\boldsymbol{h}_{2i}))$ and $\tau$ is a temperature parameter. Here, $(\boldsymbol{h}_{1i}, \boldsymbol{h}_{1j})$ are within-view negatives, and $(\boldsymbol{h}_{1i}, \boldsymbol{h}_{2j})$ are cross-view negatives.

Since the two views are symmetric, the final node-level contrastive loss is

$$\mathcal{L}_{\text{node}} = \frac{1}{2N} \sum_{i=1}^{N} \big[\ell(\boldsymbol{h}_{1i}, \boldsymbol{h}_{2i}) + \ell(\boldsymbol{h}_{2i}, \boldsymbol{h}_{1i})\big]. \tag{5}$$

Minimizing this loss pulls positive pairs together while pushing negative pairs apart, preserving biological variability across cells and implicitly correcting batch effects, ultimately yielding more robust node embeddings.

##### A.4 Alternating topology–attribute denoising.

Single-cell proteomic data contain substantial noise from sample preparation and MS quantification. To obtain noise-resilient embeddings, we adopt an alternating denoising strategy consisting of (i) attribute denoising and (ii) topology-based contrastive learning.

**Attribute denoising.** To reduce noise in the proteomic profiles, we use prototype contrastive learning. Prototypes lying far from cluster boundaries are more robust to noise [6], so we leverage them to guide embedding refinement. At each epoch, we cluster the current embeddings (without augmentation) into $K$ groups via $k$-means, yielding cluster assignments $\text{cluster}_i$ and cluster centers $\{c_1, \ldots, c_K\}$. Each prototype $c_k$ is the mean embedding of nodes in cluster $k$:

$$c_k = \frac{1}{\text{Num}_k} \sum_{\text{cluster}_i = k} \boldsymbol{h}_i, \tag{6}$$

where $\text{Num}_k$ is the number of nodes assigned to cluster $k$.

We then update all embeddings using the prototype-level contrastive loss

$$\mathcal{L}_{\text{proto}} = -\frac{1}{N} \sum_{i=1}^{N} \log \frac{\exp(\boldsymbol{h}_i \cdot c_{\text{cluster}_i}/\tau)}{\sum_{k=1}^{K} \exp(\boldsymbol{h}_i \cdot c_k/\tau)}. \tag{7}$$

**Overall loss.** The final objective combines node-level contrastive learning (based on the augmented embeddings $\boldsymbol{H}_1$ and $\boldsymbol{H}_2$) with prototype denoising:

$$\mathcal{L} = \mathcal{L}_{\text{node}} + \gamma\, \mathcal{L}_{\text{proto}}, \tag{8}$$

where $\gamma$ is a balancing coefficient (set to 0.05 in our experiments). The model is optimized using Adam with a learning rate of $10^{-3}$.

### B Evaluation Metrics

The metrics are formally defined as follows:

**Adjusted Rand Index (ARI):**

$$\text{ARI} = \frac{\sum_{ij}\binom{n_{ij}}{2} - \frac{\sum_i \binom{a_i}{2}\sum_j \binom{b_j}{2}}{\binom{n}{2}}}{\frac{1}{2}\left[\sum_i \binom{a_i}{2} + \sum_j \binom{b_j}{2}\right] - \frac{\sum_i \binom{a_i}{2}\sum_j \binom{b_j}{2}}{\binom{n}{2}}}$$

where $n_{ij}$ denotes the number of cells assigned to both true cluster $i$ and predicted cluster $j$, and $a_i$, $b_j$ represent the respective cluster marginals.

**Average Silhouette Width (ASW):**

$$\text{ASW} = \frac{1}{N}\sum_i \frac{p(i) - q(i)}{\max\{p(i), q(i)\}}$$

where $q(i)$ is the average intra-cluster distance for cell $i$, $p(i)$ is the smallest average inter-cluster distance, and $N$ is the total number of cells.

**Normalized Mutual Information (NMI):**

$$\text{NMI} = \frac{MI(\hat{Y};Y)}{\sqrt{H(\hat{Y})H(Y)}}$$

where $MI(\hat{Y};Y)$ denotes the mutual information between predicted cluster assignments $\hat{Y}$ and true labels $Y$, and $H(\cdot)$ represents Shannon entropy.

**Purity Score (PS):**

$$\text{PS} = \frac{1}{N}\sum_i \max_j n_{ij}$$

which measures the proportion of correctly assigned cells per predicted cluster.

These complementary metrics jointly evaluate the accuracy, separability, consistency, and compositional purity of the resulting cell clusters, providing a comprehensive assessment of clustering performance.

### C Additional Experiments

#### C.1 t-SNE Results

The six t-SNE panels provide a qualitative comparison of cell embeddings produced by different encoder types on the integrated single-cell proteomics datasets of N2 and nanoPOTS. The GCN encoder shows moderate separation between some cell types, but clusters are diffuse and partially overlapping, with uneven batch mixing. The Beta Kernel Direct encoder yields compact, well-defined clusters with substantially improved batch mixing, suggesting strong alignment and biological signal preservation. The RWR-Truncated encoder produces scattered clusters with elongated or split shapes and inconsistent mixing, indicating an underrepresentation of the global topology. The RWR-Interpolated encoder improves cluster coherence and dataset alignment relative to the truncated version, better preserving cell identities. The Heat Kernel-Approximated encoder generates compact clusters with good local separation, though some overlap remains, while the Heat Kernel-Truncated encoder shows tight, distinct clusters with robust batch harmonization. Overall, the Beta Kernel Direct and Heat Kernel-Truncated encoders visually perform best, RWR-Interpolated and Heat Kernel-Approximated show moderate performance, and GCN and RWR-Truncated are the weakest. These results highlight that spectral- and kernel-based encoders generally produce more biologically meaningful embeddings than baseline GCNs or truncated proximity encoders.

#### C.2 Effect of Polynomial Degree

In Fig. 1, we evaluate the clustering performance of the interpolated Heat Kernel, interpolated RWR kernel, and the direct Beta Kernel across varying polynomial degrees $K$ on the Scope2_Specht dataset. Across all four evaluation metrics—ARI, ASW, NMI, and PS—the interpolated Heat Kernel generally exhibits strong and stable behavior for moderate values of $K$, while the interpolated RWR kernel shows a gradual improvement, particularly at higher polynomial degrees. The direct Beta Kernel remains consistently competitive, achieving robust performance without requiring interpolation. When compared against the non-spectral baseline GCN Encoder, all three kernel-based methods demonstrate comparable or superior results, especially in terms of ASW and PS. Overall, the figure highlights the advantages of Arnoldi-orthonormalization-based spectral GCN approach, showing that increasing the polynomial degree can improve clustering stability and quality, particularly for the interpolated RWR-based representations.

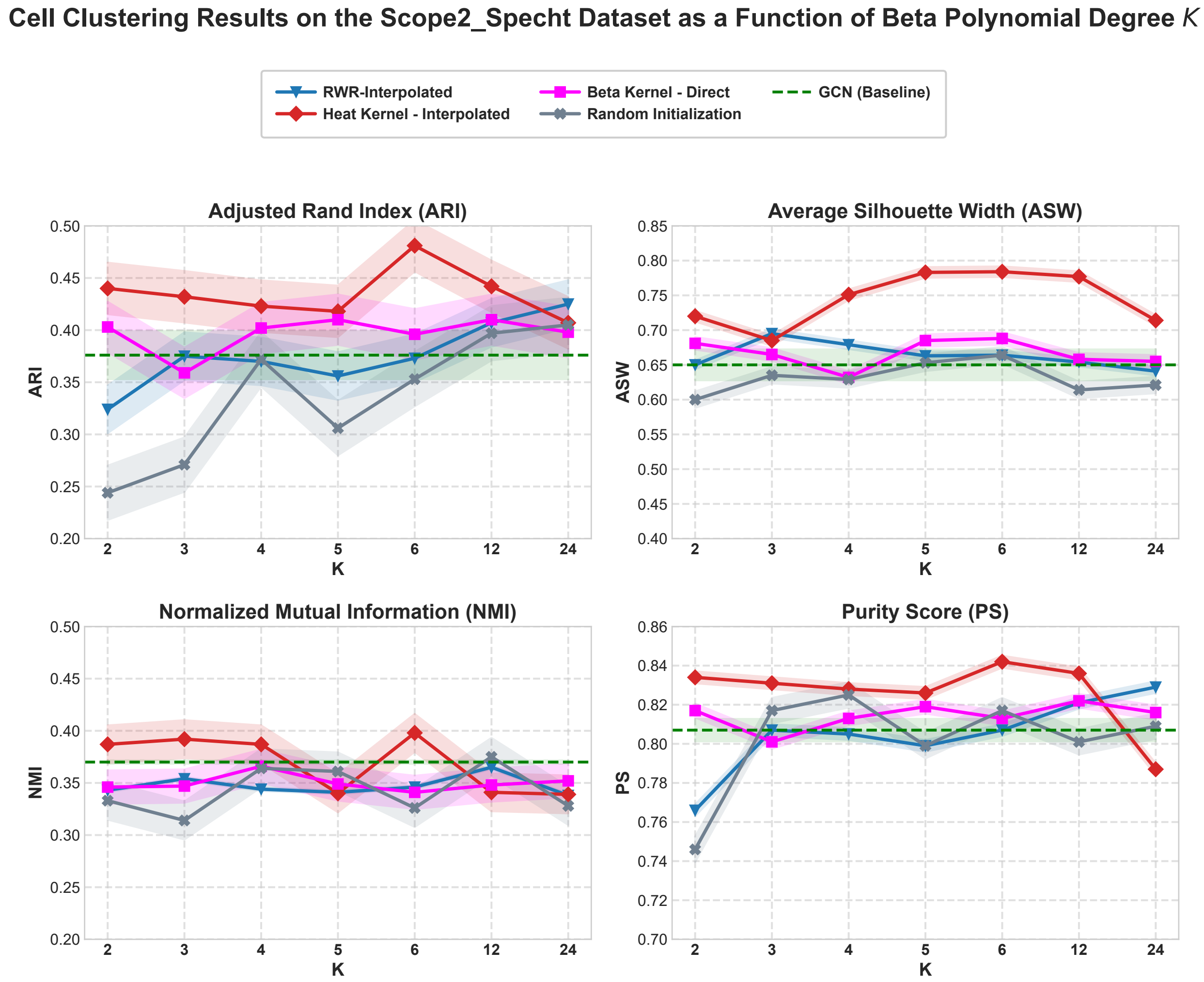


Fig. 1: Comparison of cell clustering performance on the `Scope2_Specht` dataset using Beta Kernel, Heat Kernel (Interpolated), and RWR-Interpolated filters across varying polynomial degrees $K \in \{2, 3, 4, 5, 6, 12, 24\}$. The four evaluation metrics include ARI,ASW, NMI, and PS. non-spectral GCN-Encoder is shown for baseline reference. The results illustrate how spectral interpolation and polynomial degree influence clustering performance across multiple criteria. The figure shows Mean ± 95% Confidence Interval over eight different seed runs.

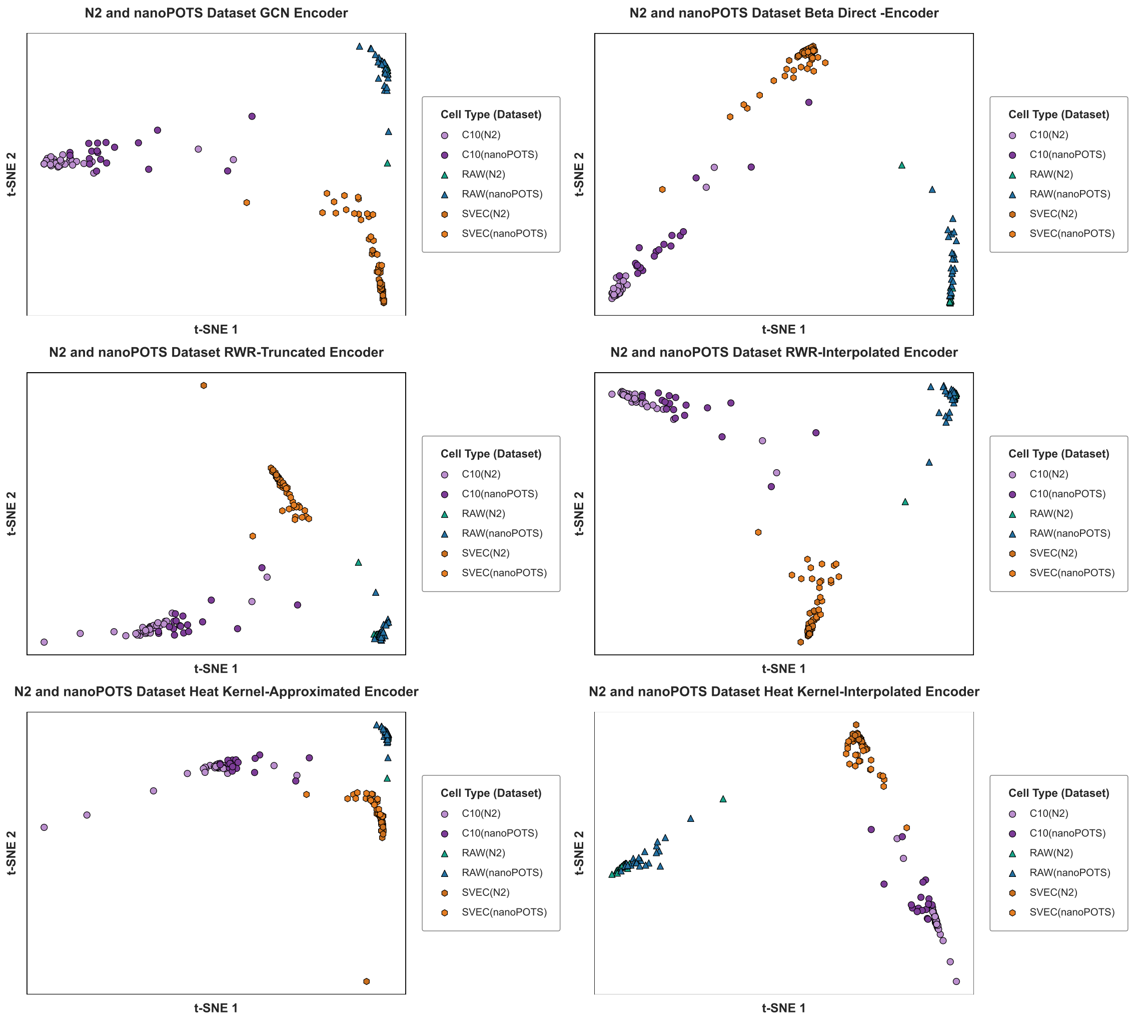


Fig. 2: t-SNE visualizations of cells from the N2 and nanoPOTS datasets. Each plot shows cells colored by their data acquisition batch and cell type. The six panels correspond to the different encoders: (i) GCN Encoder, (ii) Beta Kernel Direct, (iii) RWR Truncated, (iv) RWR Interpolated, (v) Heat Kernel Approximated, and (vi) Heat Kernel Truncated.

#### C.3 Effect of Embedding Dimension

To assess how the embedding dimension $D$ influences clustering performance, we evaluated multiple metrics—Adjusted Rand Index (ARI), Average Silhouette Width (ASW), Normalized Mutual Information (NMI), and Purity Score (PS)—across six encoder types on the Scope2_Specht dataset (Figure 3).

Overall, moderate dimensions ($D = 64$) tend to yield the best performance for most encoders. The GCN encoder achieves its highest ARI and NMI at $D = 64$, while larger dimensions do not consistently improve clustering quality and sometimes reduce cluster coherence. Beta Kernel Direct embeddings exhibit relatively stable performance across dimensions, with peak ARI at $D = 64$ and minimal variation in ASW and PS. Heat Kernel Interpolated embeddings also favor $D = 64$, showing improved ARI, ASW, and NMI, whereas the Heat Kernel Approximated encoder demonstrates slightly declining performance as $D$ increases beyond 64. For RWR-based encoders, the RWR-Interpolated method performs best at moderate dimensions, while RWR-Truncated shows a delayed peak at $D = 256$, reflecting sensitivity to global topological information.

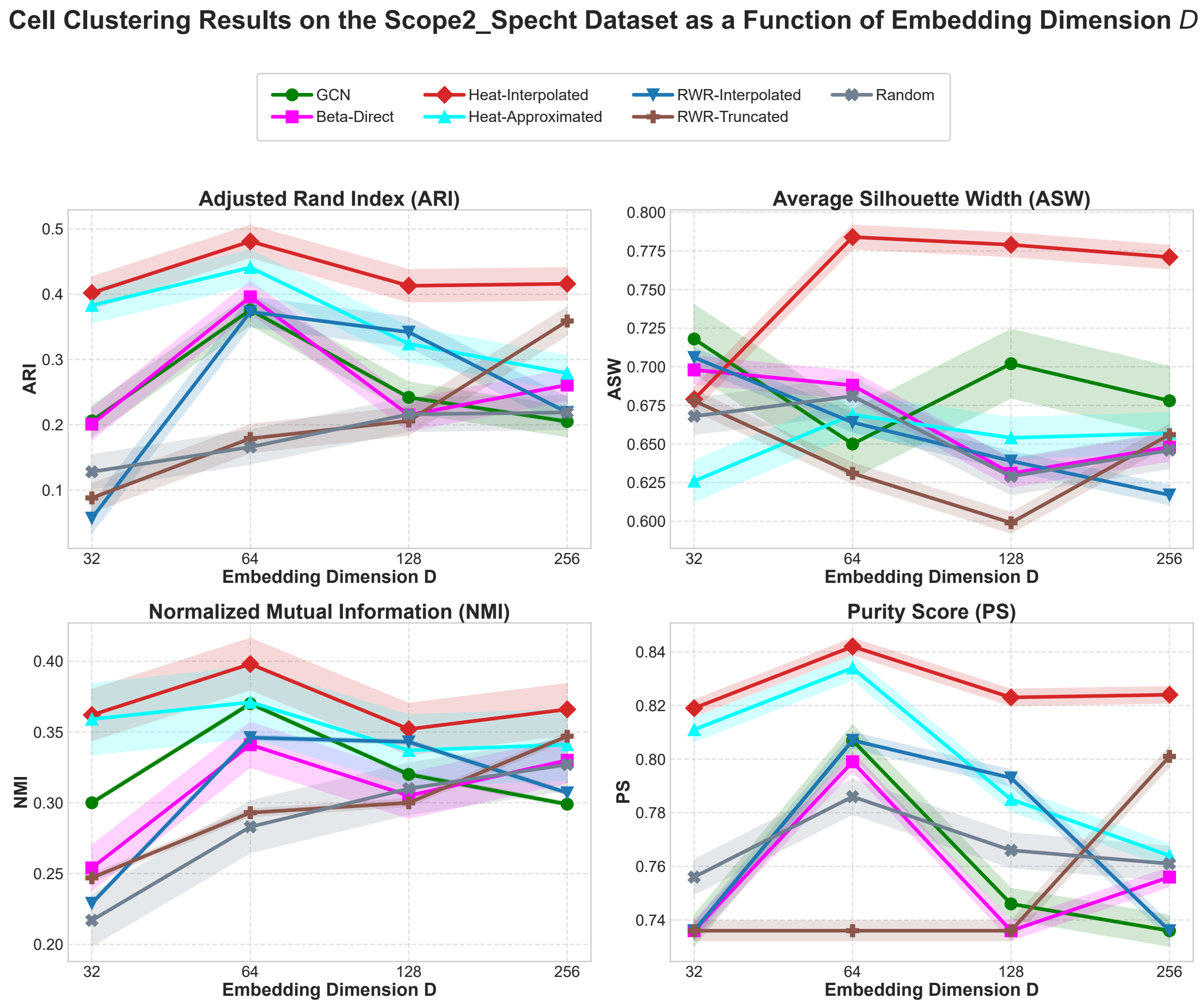


Fig. 3: Clustering performance as a function of the embedding dimension $D \in \{32, 64, 128, 256\}$ for six graph-based representation learning methods. The evaluation includes ARI, ASW, NMI, and PS metrics, with the x-axis shown on a logarithmic scale to reflect the exponential growth of $D$. The figure shows Mean ± 95% Confidence Interval over eight different seed runs.

Table 1: Summary of hyperparameters used in the experiments.

| Hyperparameter | Value | Description |
|---|---|---|
| `stage1` | True | Start from Stage 1 preprocessing |
| Learning rate | $1 \times 10^{-3}$ | Optimization step size |
| Hidden dimension | 64 | GNN latent dimension |
| Projection head dim. | 256 | Dimension of projection layer |
| Activation | ReLU | Nonlinear activation function |
| # GCN layers | 3 | Depth of encoder network |
| # Prototypes | 2 | Structural prototype count |
| Perturbed edges | 50 | Added/removed edges for augmentation |
| Topology denoising | False | Enable structural denoising |
| DropEdge rate (view 1) | 0.2 | Edge dropout for view 1 |
| DropEdge rate (view 2) | 0.4 | Edge dropout for view 2 |
| Feature mask (view 1) | 0.4 | Feature dropout for view 1 |
| Feature mask (view 2) | 0.2 | Feature dropout for view 2 |
| $\alpha$ | 0.05 | Weighting factor in contrastive loss |
| $\tau$ | 0.4 | Temperature parameter |
| Weight decay | $1 \times 10^{-5}$ | $L_2$ regularization |
| Epochs | 200 | Number of training epochs |
| Random seed | 39788 | Reproducibility seed |
| Graph threshold | 0.15 | Similarity cutoff for edges |
| Feature preprocessing | True | Apply feature normalization |

##### C.4 Hyperparameters used in the experiments

To ensure stable and reproducible training, we follow a consistent set of hyperparameters across all experiments. The framework begins from `Stage 1` by default and applies feature preprocessing before graph construction using a similarity threshold of 0.15. We train the encoder for 200 epochs with a learning rate of $10^{-3}$, weight decay of $10^{-5}$, and a random seed of 39788. The GNN backbone uses a hidden dimension of 64, six propagation layers, and ReLU activations. For contrastive learning, we adopt a projection head of dimension 256, two structural prototypes, and a balance coefficient $\alpha = 0.05$. Data augmentations include edge dropping (0.2/0.4 for the two graph views) and feature masking (0.4/0.2). Additional parameters include a temperature $\tau = 0.4$, a perturbation budget of 50 edges, and an optional topology-denoising module, disabled by default. All experiments are performed on Google Colab with NVIDIA A100-SXM4 GPU with 40 GB of HBM2 VRAM, complemented by 83.5 GB of system RAM and 235.7 GB of available disk storage..

##### C.5 Effect of Graph Edge Cut Threshold

To study the influence of the graph edge cut threshold $h$ on clustering performance, we evaluate multiple metrics—Adjusted Rand Index (ARI), Average Silhouette Width (ASW), Normalized Mutual Information (NMI), and Purity Score (PS)—across different embedding and propagation methods on the Scope2_Specht dataset (Figure **??**). In our graph construction procedure, we first compute a cellwise similarity matrix $S$ using the Pearson correlation coefficient (PCC) between the abundance feature vectors of cells, i.e.,

$$S_{ij} = \mathrm{PCC}(v_i, v_j) = \frac{\mathrm{cov}(x_i, x_j)}{\sigma_{x_i}\sigma_{x_j}},$$

where $\mathrm{cov}(\cdot,\cdot)$ and $\sigma$ denote the covariance and standard deviation, respectively. Then, a threshold $h$ is applied to obtain the adjacency matrix $A \in \mathbb{R}^{N\times N}$ of the cell graph, defined as

$$A_{ij} = \begin{cases} 1, & \text{if } S_{ij} > h, \\ 0, & \text{otherwise.} \end{cases}$$

In this setting, the edge cut threshold $h$ directly controls the sparsity of the constructed graph: smaller values of $h$ yield denser graphs, while larger values result in sparser topologies.

Table 2: **Runtime of GCL algorithms and spectral encoders on the** `Scope2_Specht` **data.**

| Cut-off Value | Step | GCN | Heat Kernel | Random Walk | Beta Kernel |
|---|---|---|---|---|---|
| **h= 0.05** | Phase 1 training | 1937.25s | | | |
| | Graph construction | 36.71s ± 1.88s | | | |
| | Phase 2 Training | 19.32s ± 0.10s | 19.52s ± 0.11s | 19.43s ± 0.10s | 19.14s ± 0.87s |
| | Interpolation (K=12) | None | 0.000817s ± 0.000014s | 0.000793s ± 0.000016s | None |
| | Encoder Training | 0.00487s ± 0.00015s | K=3 0.00568s ± 0.00011s | K=3 0.00609s ± 0.00036s | K=3 0.00568s ± 0.00071s |
| | | | K=6 0.00639s ± 0.00016s | K=6 0.00614s ± 0.00005s | K=6 0.00559s ± 0.00030s |
| | | | K=12 0.00846s ± 0.00046s | K=12 0.00767s ± 0.00003s | K=12 0.00573s ± 0.00007s |
| **h= 0.3** | Phase 1 training | Retained | | | |
| | Graph construction | 29.90s ± 0.44s | | | |
| | Phase 2 Training | 7.82s ± 0.05s | 8.11s ± 0.06s | 8.03s ± 0.07s | 8.18s ± 0.04s |
| | Interpolation (K=12) | None | Retained | Retained | None |
| | Encoder Training | 0.00466s ± 0.00006s | K=3 0.00543s ± 0.00010s | K=3 0.00544s ± 0.00005s | K=3 0.00518s ± 0.00012s |
| | | | K=6 0.00639s ± 0.00016s | K=6 0.00605s ± 0.00008s | K=6 0.00529s ± 0.00015s |
| | | | K=12 0.00774s ± 0.00014s | K=12 0.00756s ± 0.00005s | K=12 0.00565s ± 0.00019s |
| **h= 0.75** | Phase 1 training | Retained | | | |
| | Graph construction | 28.85s ± 0.33s | | | |
| | Phase 2 Training | 7.20s ± 0.07s | 8.00s ± 0.09s | 8.34s ± 0.06s | 8.20s ± 0.03s |
| | Interpolation (K=12) | None | Retained | Retained | None |
| | Encoder Training | 0.00684s ± 0.00006s | K=3 0.00550s ± 0.00006s | K=3 0.00541s ± 0.00004s | K=3 0.00539s ± 0.00034s |
| | | | K=6 0.00598s ± 0.00009s | K=6 0.00610s ± 0.00004s | K=6 0.00538s ± 0.00007s |
| | | | K=12 0.00771s ± 0.00002s | K=12 0.00749s ± 0.00005s | K=12 0.00568s ± 0.00012s |

##### C.6 Runtime Analysis

The runtime analysis in Table 2 shows clear computational differences among the methods. GCN-based approaches are consistently the fastest due to their localized message-passing mechanism and reliance on sparse matrix operations, which scale efficiently with the number of edges. In contrast, Heat- and RWR-based methods incur higher computational cost because they involve diffusion processes, iterative propagation, and polynomial or spectral approximations that require additional matrix computations. Moreover, the observed decrease in runtime as the threshold $h$ increases can be attributed to the induced sparsity of the graph, which reduces the number of active edges and thus the overall computational burden. The small confidence intervals further indicate stable and consistent runtime behavior across runs.
